# An emergent population code in primary auditory cortex supports selective attention to spectral and temporal sound features

**DOI:** 10.1101/2020.03.09.984773

**Authors:** Joshua D. Downer, Jessica R. Verhein, Brittany C. Rapone, Kevin N. O’Connor, Mitchell L. Sutter

## Abstract

Textbook descriptions of primary sensory cortex (PSC) revolve around single neurons’ representation of low-dimensional sensory features, such as visual object orientation in V1, location of somatic touch in S1, and sound frequency in A1. Typically, studies of PSC measure neurons’ responses along few (1 or 2) stimulus and/or behavioral dimensions. However, real-world stimuli usually vary along many feature dimensions and behavioral demands change constantly. In order to illuminate how A1 supports flexible perception in rich acoustic environments, we recorded from A1 neurons while rhesus macaques performed a feature-selective attention task. We presented sounds that varied along spectral and temporal feature dimensions (carrier bandwidth and temporal envelope, respectively). Within a block, subjects attended to one feature of the sound in a selective change detection task. We find that single neurons tend to be high-dimensional, in that they exhibit substantial mixed selectivity for both sound features, as well as task context. Contrary to common findings in many previous experiments, attention does not enhance the single-neuron representation of attended features in our data. However, a population-level analysis reveals that ensembles of neurons exhibit enhanced encoding of attended sound features, and this population code tracks subjects’ performance. Importantly, surrogate neural populations with intact single-neuron tuning but shuffled higher-order correlations among neurons failed to yield attention-related effects observed in the intact data. These results suggest that an emergent population code not measurable at the single-neuron level might constitute the functional unit of sensory representation in PSC.

**SIGNIFICANCE STATEMENT:** The ability to adapt to a dynamic sensory environment promotes a range of important natural behaviors. We recorded from single neurons in monkey primary auditory cortex while subjects attended to either the spectral or temporal features of complex sounds. Surprisingly, we find no average increase in responsiveness to, or encoding of, the attended feature across single neurons. However, when we pool the activity of the sampled neurons via targeted dimensionality reduction, we find enhanced population-level representation of the attended feature and suppression of the distractor feature. This dissociation of the effects of attention at the level of single neurons vs. the population highlights the synergistic nature of cortical sound encoding and enriches our understanding of sensory cortical function.

## INTRODUCTION

Classic accounts of primary sensory cortex (PSC) relegate PSC function to sensory filtering. Accordingly, individual PSC neurons act as independent filters for low-dimensional sensory features (Hubel and Wiesel, 1968; Kaas et al., 1979; Merzenich et al., 1975) while “association” and prefrontal cortical neurons integrate information about behavioral demands and other sensory modalities (Robinson et al., 1978). This account supports a feed-forward nervous system model, where information propagates along distinct processing stages, from peripheral sensory receptors to motor effectors, to enable perception and behavior (Riesenhuber and Poggio, 1999; Van Essen and Maunsell, 1983). A wealth of studies has shown that PSC neurons often form maps of the sensory epithelium, consistent with a role as low dimensional filters (e.g. (Merzenich et al., 1975)). Moreover, secondary and higher-order cortical neurons exhibit diminished, more complex, and task-dependent versions of these maps (Rauschecker and Scott, 2009), and prefrontal cortical neurons seem to represent all manner of stimulus and cognition-related variables without clear functional topography (Machens et al., 2010).

Primary auditory cortex (A1) receives significant feed-forward input from the lemniscal auditory thalamus and is therefore understood as the earliest cortical auditory processing stage. However, decades of research confound categorization of A1 neurons as static sensory filters: myriad non-sensory variables affect the firing rates, variance and noise correlations in A1 (David, 2018; Osmanski and Wang, 2015).

Generally, these studies have shown that, under demanding sensory conditions, A1 neuron responses and/or tuning are enhanced, consistent with findings across PSC modalities (Carlson et al., 2018; Gomez-Ramirez et al., 2016; Mineault et al., 2016). Thus, a widely held view is that A1 and other PSCs are best described as arrays of adaptive sensory filters, whereby simple feature tuning is modulated by ongoing behavioral and sensory demands. However, recent studies show that A1 neurons’ synergistic interactions can play a large role in sensory processing – a role which often cannot be understood solely based on the activity of individual neurons (Bagur et al., 2018; Bathellier et al., 2012; Harris et al., 2011; See et al., 2018). Rather, individual neurons’ activity may be better understood in terms of their contributions within functional ensembles.

We recorded from A1 neurons while monkeys performed a task wherein attention is switched between two different features of a sound (Figure 1). We find that A1 neurons robustly represent each task variable: both sound features, as well as task context. Importantly, contrary to a vast literature across sensory modalities showing that attention improves single-neuron sensory encoding, we find no overall or average attentional modulation of single A1 neurons. The fact that A1 neurons encode each sound feature as well as task context but display no average attention-related sensory enhancement suggests that A1 single-neuron activity in this task is not directly related to task performance. These null effects surprised us, since the task presents significant auditory sensory demands related to sound features encoded by A1 neurons. This prompted us to conduct a population-level analysis, inspired by studies performed in PFC (Mante et al., 2013; Rigotti et al., 2013), to make sense of the heterogeneous single-neuron representations.

**Figure 1.**
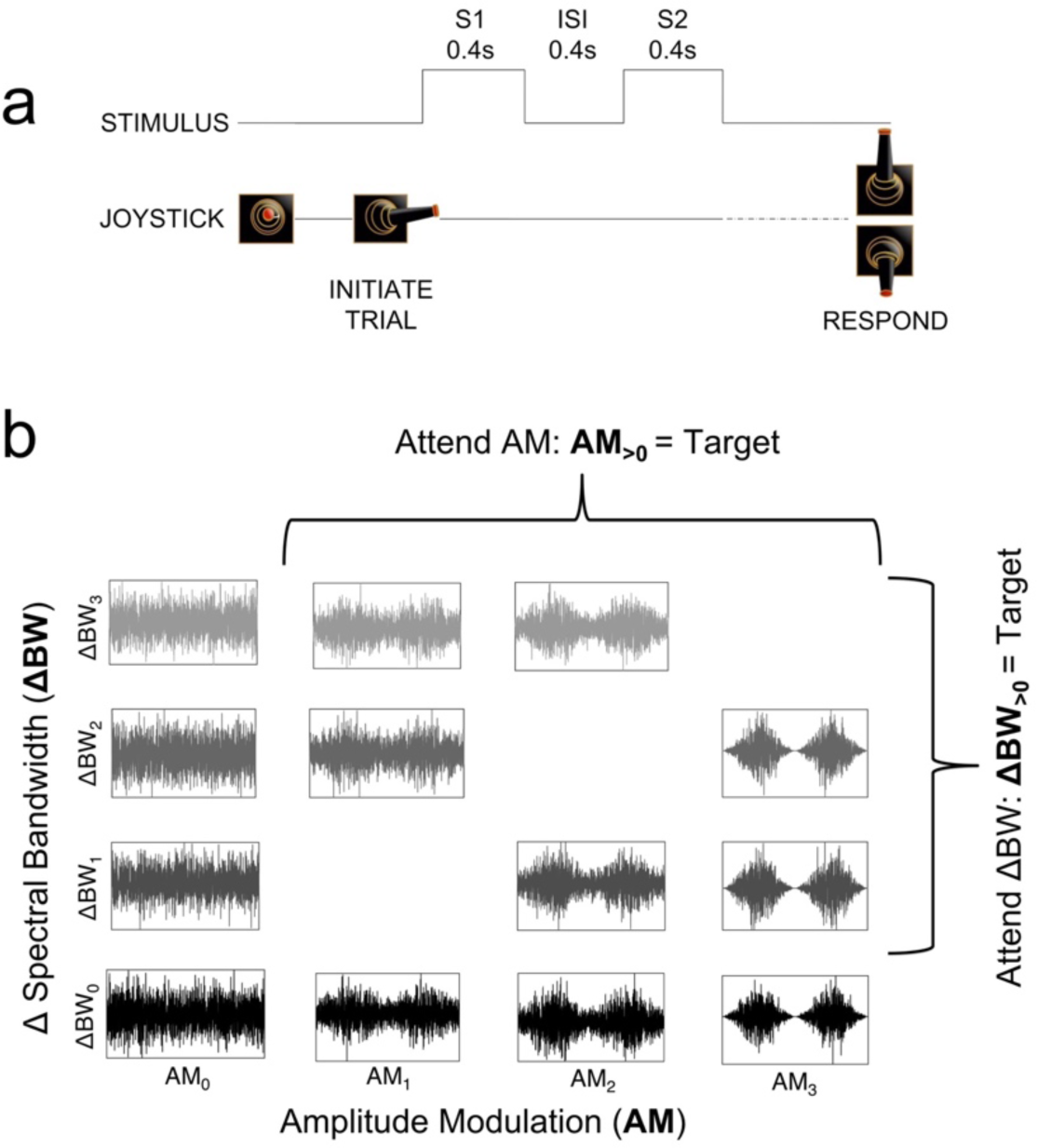
Task and stimulus design. (**a**) Subjects initiated a trial by moving a joystick laterally to present 2 sequential 0.4-s sounds (S1 and S2) separated by a 0.4-s silent inter-sound interval (ISI). Subjects then used the joystick to make a behavioral report, a “Yes” or “No” response indicating whether they detected the attended sound feature in S2. (**b**) Sounds varied along spectral (bandwidth, ΔBW) and temporal (amplitude modulation, AM) feature dimensions. Subjects were trained to selectively attend to changes in one feature dimension or the other across blocks. The S1 sound was always unmodulated and full bandwidth (AM0-ΔBW0, bottom left) and the S2 could be any sound in the set. During the Attend AM condition all sounds with amplitude modulation (i.e., with AM1, AM2, or AM3) were targets, and during the Attend BW condition all sounds with narrower spectral bandwidth than the standard (ΔBW1, ΔBW2, and ΔBW3) were targets.

Our population analyses focused on reducing the dimensionality of population activity from the number of neurons to the number of variables. Thus, we defined a low-dimensional subspace in which the stimulus and behavioral variables are encoded at the population level (Gao et al., 2017). Neural activity projected into this subspace revealed strong effects of attention on population-level sound feature encoding, where the population sensitivity to the attended feature is enhanced. Moreover, variability in these projections accounts for subjects’ performance. Further analyses using “surrogate” populations for comparison, in which the single neuron marginal statistics are kept intact while shuffling higher order interactions, (Elsayed and Cunningham, 2017), reveal that our findings don’t arise as an expected by-product of pooling across many neurons. Rather, these results suggest that sound-encoding in A1 relies on synergies among neurons, and these synergies support the neural code for selective listening.

## MATERIALS AND METHODS

### Subjects

We recorded from 92 A1 neurons of two adult rhesus macaques, one female (W, 8 kg) and one male (U, 12 kg). Subjects were each implanted with a head fixation post and a recording cylinder over a left-sided 18mm parietal craniotomy. Craniotomies were centered over auditory cortex as determined by stereotactic coordinates, allowing for vertical electrode access to the superior temporal plane via parietal cortex (Saleem and Logothetis, 2012). Surgical procedures were performed under aseptic conditions; all animal procedures met the requirements set forth in the United States Public Health Service Policy on Humane Care and Use of Experimental Animals and were approved by the Institutional Animal Care and Use Committee of the University of California, Davis.

### Stimuli

Sound stimuli varied along spectral and temporal dimensions (either or both; Figure 1b). S1 was always unmodulated, full-bandwidth Gaussian white noise with a 9 octave range (40 to 20,480 Hz; Figure 1b, bottom left in stimulus grid: AM0ΔBW0). Noise signals were generated from four seeds, which were frozen across recording sessions. To introduce spectral and temporal variance, the S1 sound was bandpass filtered to narrow the spectral bandwidth (ΔBW; range: 0.375-1.5 octaves less than the 9-octave full BW) and/or sinusoidally amplitude modulated (AM; range: 28-100% of the depth of the original). Our bandpass filtering method relied on sequential single-frequency addition, thereby reducing envelope variations produced by other filtering procedures (Strickland and Viemeister, 1997). S2 could be any of the stimuli in the set represented in the grid in Figure 1b.

The precise AM and ΔBW values used during recording sessions were tailored to each subject’s psychophysical thresholds for detection of each sound feature (AM and ΔBW) in isolation, determined prior to recording. Three values of each feature were used in recording sessions: one near psychophysical threshold (defined as the modulation level at which the subject’s sensitivity, measured using *d*’ (Wickens, 2002) was equal to 1), one slightly above threshold (*d*’ ∼ 1.2), and one well above threshold (*d*’> 1.5). Experimental values of ΔBW were 0.375, 0.5, and 1 octave for monkey U and 0.5, 0.75, and 1.5 octaves for monkey W. AM depth values were 28, 40, and 100% for monkey U and 40, 60, and 100% for monkey W.

In analyses in which we collapse data across monkeys, modulation values are presented as ranks relative to subjects’ thresholds (AM0-3 and ΔBW0-3, where ‘0’ in the subscript reflects no change in that feature dimension relative to the unmodulated, full-bandwidth; Figure 1b). To reduce the size of the stimulus space, we presented 13 total stimuli during each experimental session, using only a subset of the possible comodulated stimuli (sounds with both AM and ΔBW; Figure 1B). The AM frequency was held constant in each individual session but varied from day to day. Across sessions we randomly selected the AM frequency from a small range of frequencies (15, 22, 30, 48, and 60 Hz). Previous work has shown that rhesus macaques’ average AM detection thresholds are similar across the full range of frequencies used in our task (O’Connor et al., 2011). We chose to randomly select the presented AM frequency to avoid biasing our recordings toward AM-sensitive neurons at the expense of sampling from ΔBW-sensitive neurons.

Sounds were 400 ms in duration with 5-ms cosine ramps at onset and offset. Sounds were presented from a single speaker (RadioShack PA-110 or Optimus Pro-7AV) positioned approximately 1m in front of the subject, centered interaurally. Each stimulus was calibrated to an intensity of 65 dB SPL at the outer ear (A-weighted; Bruel & Kjaer model 2231).

### Feature selective attention task

Subjects sat in an acoustically transparent primate chair and used a joystick to perform a ‘Yes-No’ task (Figure 1a). LED illumination cued the onset of each trial.

Subjects initiated sound presentation with a lateral joystick movement, which was followed by a 100-ms delay and two sequential sounds (S1 then S2) separated by a 400-ms inter-sound interval (ISI). The trial was aborted if the lateral joystick position was not maintained for the full duration of both stimuli, and an interrupt timeout (5-10 s) was enforced. The first stimulus (S1) was always the unmodulated, full-bandwidth (9-octaves) Gaussian white noise. The second stimulus (S2) was chosen from the set in Figure 1B. S2 sounds were presented pseudorandomly, such that the entire stimulus set across all four noise seeds was represented over each set of 96 trials.

The target feature (ΔBW or AM) alternated by block. Subjects were given an LED cue (green or red, counterbalanced across subjects) for which feature to attend, positioned above the speaker. The first 60-180 trials of each block served as “instruction trials,” containing the S1 and sounds with a single target feature (e.g., for an attend AM block all stimuli in the instruction block were full-bandwidth (ΔBW0) with AM as the variable (AM0-3), This allowed the animal to more reliably focus on the attended feature without the distraction of variation in the unattended feature. The length of instruction blocks depended on subjects’ performance: if 78% of trials were not correctly performed within 60 trials, another instruction block was begun, up to a limit of 3 instruction blocks (180 trials). Subsequently an attention block was begun. After the offset of S2, subjects were required to respond with a “yes” joystick movement on trials in which the target feature was present ([Fig. 1b]: AM1-3 during Attend AM and ΔBW1-3 during Attend BW). “No” responses were made by moving the joystick in the opposite direction. The direction to move the joystick for “yes” and “no” responses was counterbalanced between subjects. Subjects were rewarded for hits and correct rejections with water or juice and penalized with a white-light-cued timeout (5-10 s) for false alarms and misses.

Block length varied from 120 to 360 trials, excluding instruction trials. Length depended in part on subjects’ performance: if and only if 78% correct was achieved in two successive 60-trial sub-blocks could the subject transition to the next attention condition. More than one block per attention condition was performed in a given session. Sessions with fewer than 180 completed trials in either attention condition were excluded from analysis.

We analyzed subjects’ average performance within each session by calculating a coefficient that quantifies the influence of each feature on subjects’ perceptual judgment. The coefficients are derived from the following binomial logistic regression model:

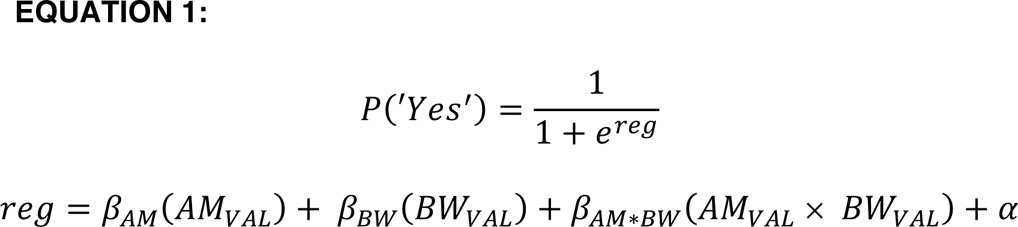

where *AM*_*VAL*_ and *BW*_*VAL*_ are the ranked values of AM and ΔBW (0 – 3), respectively, and *β* are coefficients for the value terms. We include the interaction term (β_*AM*BW*_), as well as an offset term (α) to capture response bias. Intuitively, as a given feature’s influence on the probability of a subject responding “yes” increases, the value of the feature coefficient will increase; when a given feature has no impact on the behavioral response, the value of the coefficient will be ∼0. We calculated each coefficient within each attention block (*β_AM_ β_BW_* and β_*AM*BW*_). We quantified the effect of attention by comparing the distribution of both *β_AM_ and β_BW_* between attention conditions (Figure 2).

**Figure 2.**
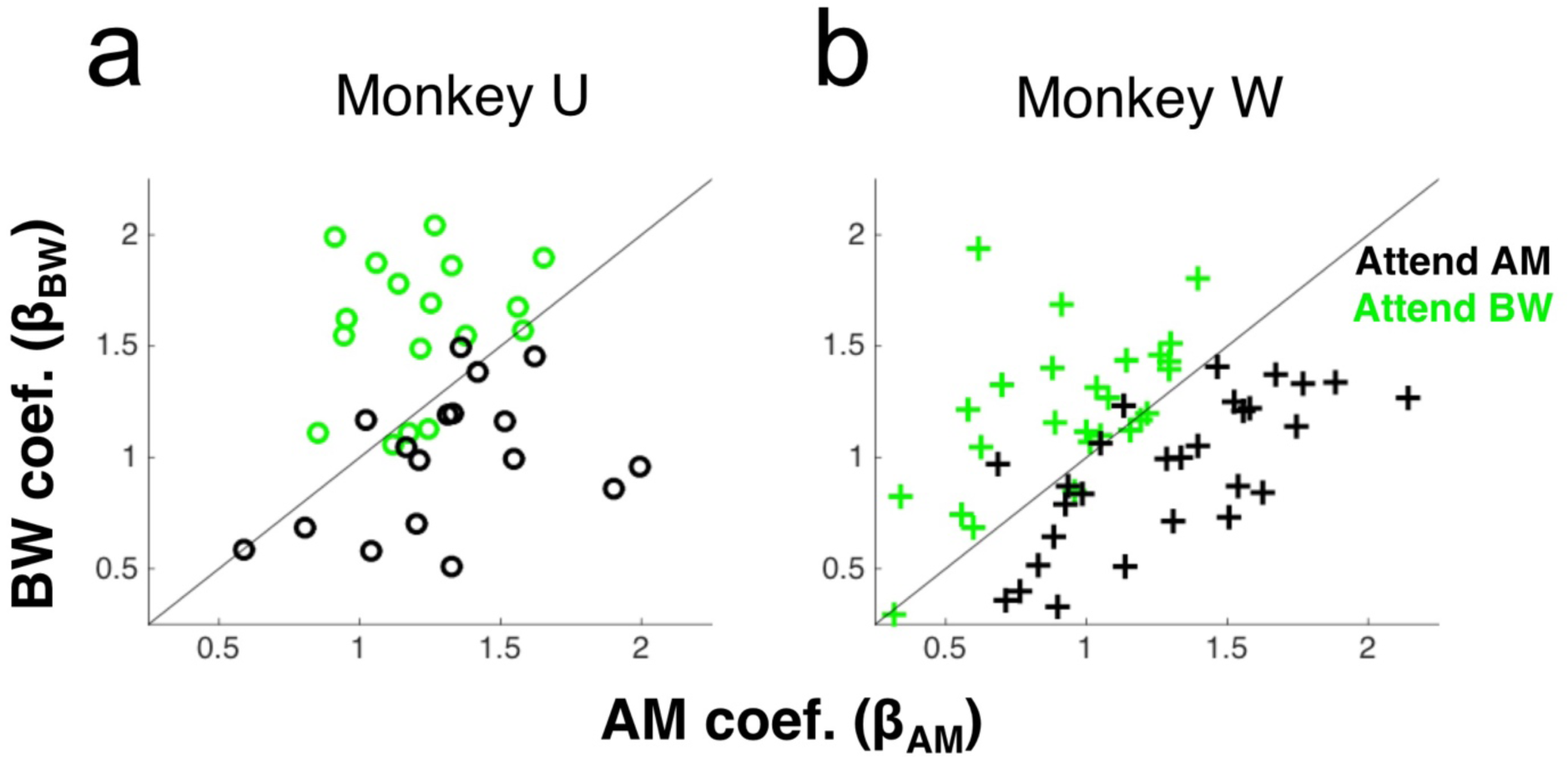
Both monkeys perform the task. (a,b) For each monkey (Monkey U in (a) and Monkey W in (b)), we show the behavioral coefficients calculated in Equation 1 across all sessions in which a recording was performed. For both subjects, ßAM is higher during the Attend AM condition than in the Attend BW condition (black markers), and vice versa (green markers). Statistical significance was determined within each monkey using a sign rank test comparing each feature’s behavioral coefficient between attention conditions (p was <0.001 for each of the 4 tests). Note that some of the sessions included in this figure were from recordings outside of A1 and therefore not included in the subsequent neural analyses.

### Single neuron electrophysiology

Recordings were performed in a sound-attenuated, foam-lined booth (IAC: 9.5′ × 10.5′ × 6.5′). We independently advanced three quartz-coated tungsten microelectrodes (Thomas Recording, 1-2 MΩ resistance, 0.35 mm horizontal spacing) in the vertical plane to the lower bank of the lateral sulcus. Pure tones and experimental stimuli were presented as the animal sat awake in the primate chair during electrode advancement, and neural responses were monitored until the multiunit activity was responsive to sound. We then attempted to isolate single units. When at least one single unit (SU) was well isolated on at least one electrode, we measured firing rate (FR) responses to at least 10 repetitions of each of the following stimuli: the unmodulated standard, ΔBW1-3 as described above, and AM sounds at 100% depth across the full range of frequencies (15, 22, 30, 48, and 60 Hz) as described above. We then cued the subject to begin the task with onset of the cue LED and recorded for the duration of the behavioral task. We repeated measurements of responses to the tested stimuli at the end of the session to ensure electrode stability. Only well isolated SUs stable over at least 120 trials for each attention condition (excluding instruction) were accepted for analysis. SU isolation was determined blind to experimental condition. SUs presented here were well isolated for a mean of 2.6 blocks, with a range of 2-5 blocks. Removing SUs with only 2 experimental blocks of isolation (n = 58) from analysis does not qualitatively affect our results nor main conclusions and interpretations.

Extracellular signals were amplified (AM Systems model 1800) and bandpass filtered between 0.3 and 10 kHz (Krohn-Hite 3382), then digitized at a 50 kHz sampling rate (Cambridge Electronic Design model 1401). Contributions of SUs to the signal were determined offline using k-means and principal component analysis-based spike sorting software in Spike2 (CED). The signal-to-noise ratio of extracted spiking activity was at least 5 standard deviations, and fewer than 0.1% of spiking events assigned to the same SU fell within a 1-ms refractory period window.

Recording locations within auditory cortical fields were estimated based on established measures of neural responses to pure-tone and bandpass noise stimuli (Petkov et al., 2006; Tian, 2001). SUs’ pure-tone frequency tuning was mapped across sessions to determine A1 boundaries based on tonotopic frequency gradients (rostral-caudal axis), width of frequency-response areas (medial-lateral axis), and response latencies. Recorded neurons were assigned to putative cortical fields post-hoc. Only SUs assigned to A1 were considered for analysis in this study.

### Analyzing task and stimulus effects in single neurons

We analyzed the effects of task context, AM and ΔBW on spike counts calculated over the entirety of each 400-ms stimulus. We used a general linear model (GLM) analysis to quantify the influences (all 3 main effects [AM, BW, and Context] as well as each of the 3 two-way interactions and the three-way interaction) of each variable on each neuron’s spike counts. We standardized spike-count (*sc*) distributions across neurons to allow for comparisons between coefficient values by *z*-scoring spike count distributions across stimuli and task context (i.e., across an entire recording). We also standardized across feature levels (which differ in their absolute values between subjects) by converting each feature value to a rank, between 0 (absence of feature) and 3 (largest feature modulation), then converting ranks to fractions varying between 0 and 1. Context was coded as a binary factor (-1 or 1 for Attend BW and Attend AM, respectively). The form of the GLM was thus:

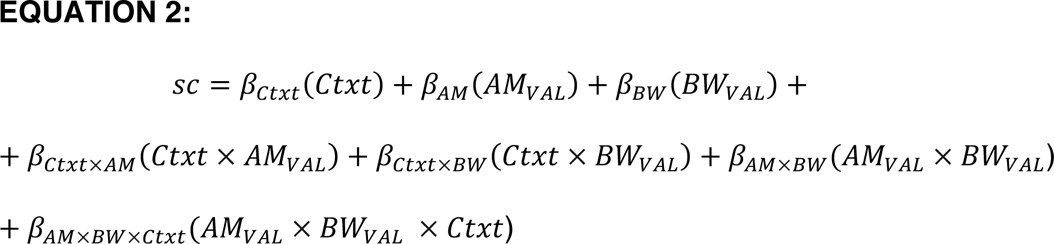

Where *sc* is the trial-by-trial spike count, *Ctxt* is the context factor, *AMVAL* and *BWVAL* are the ranked AM and BW levels, respectively and are coefficients for each factor. In a separate set of analyses, we analyzed the time course of effects using a 50-ms sliding window, between ISI onset and S2 offset. The window was shifted in 5-ms increments and we analyzed the effects of context, AM and BW and their interactions at each time point.

We also calculated the area under the Receiver Operating Characteristic (ROC area), a *sc*-based measure of sensory sensitivity that corresponds to the ability of an ideal observer to discriminate between two stimuli based on *sc* alone. Performance ranges from 0.5 (chance) to 1 (for neurons with increasing *sc* across the range of feature values) or 0 (for neurons with decreasing *sc* across the range of feature values).

ROC area can be interpreted as the probability of an ideal observer properly classifying a stimulus as containing a feature of interest, AM or ΔBW. We calculated ROC area for each stimulus condition. If a neuron’s ROC area is 0.5, that is interpreted as a failure to detect the presence of a feature. If a neuron’s ROC area is 1 or 0, that is interpreted as perfect feature detection, with either increasing (1) or decreasing (0) *sc*.

### Analyzing task and stimulus effects in neural populations

We used a “targeted dimensionality reduction” (TDR) approach to estimate the low-dimensional subspaces in which task variables may be encoded (Mante et al. 2013), and then quantified the strength of that encoding using the ROC area. TDR is a method of calculating a weighted average of neural activity across a sample of neurons. Weights for each neuron in TDR are first calculated as the linear regression coefficients relating the task variables to neurons’ firing rates (Equation 2). Then, the matrix of weights across neurons and variables is orthogonalized to allow us to estimate projections (weighted averages) of neural activity uniquely related to each task variable. TDR both reduces the dimensionality of the population response from *# neurons* to *# projections* and provides a way of “de-mixing” neural activity across a population of neurons with mixed selectivity.

TDR, and its conceptual motivation, are illustrated with a toy example in Figure 3. Here, we contrast two hypothetical cases in which feature-selective attention modulates single-neuron tuning via a feature-selective gain (FSG) mechanism (Martinez-Trujillo and Treue, 2004) by simulating two separate populations of 100 sensory neurons responding to stimuli varying along two feature dimensions. FSG is a model of attention whereby the response of a neuron is determined by both its sensory response function and a multiplicative gain parameter that depends on the neuron’s preference for the attended feature, yielding larger responses when attention is directed toward the neuron’s preferred feature. In the example case of ‘Unmixed Selectivity’ (Fig. 3a-d), single neurons’ tuning strength to either of two features (Feature A and Feature B) is largely mutually exclusive (i.e. non-zero tuning strength for Feature A is associated with ∼0 tuning strength for Feature B, and vice versa). Simulated neurons’ firing rates are thus a function of only one feature and a FSG parameter that multiplicatively scales neurons’ firing rates according to whether attention is directed toward Feature A (Fig. 3a) or Feature B (Fig. 3b). In this way, the slope of the “feature tuning strength vs. firing rate” plot increases for the attended feature (e.g., compare the red dots in Fig. 3a and b). This leads to a larger response to a feature when attention is directed toward that feature (Fig. 3c, d).

**Figure 3.**
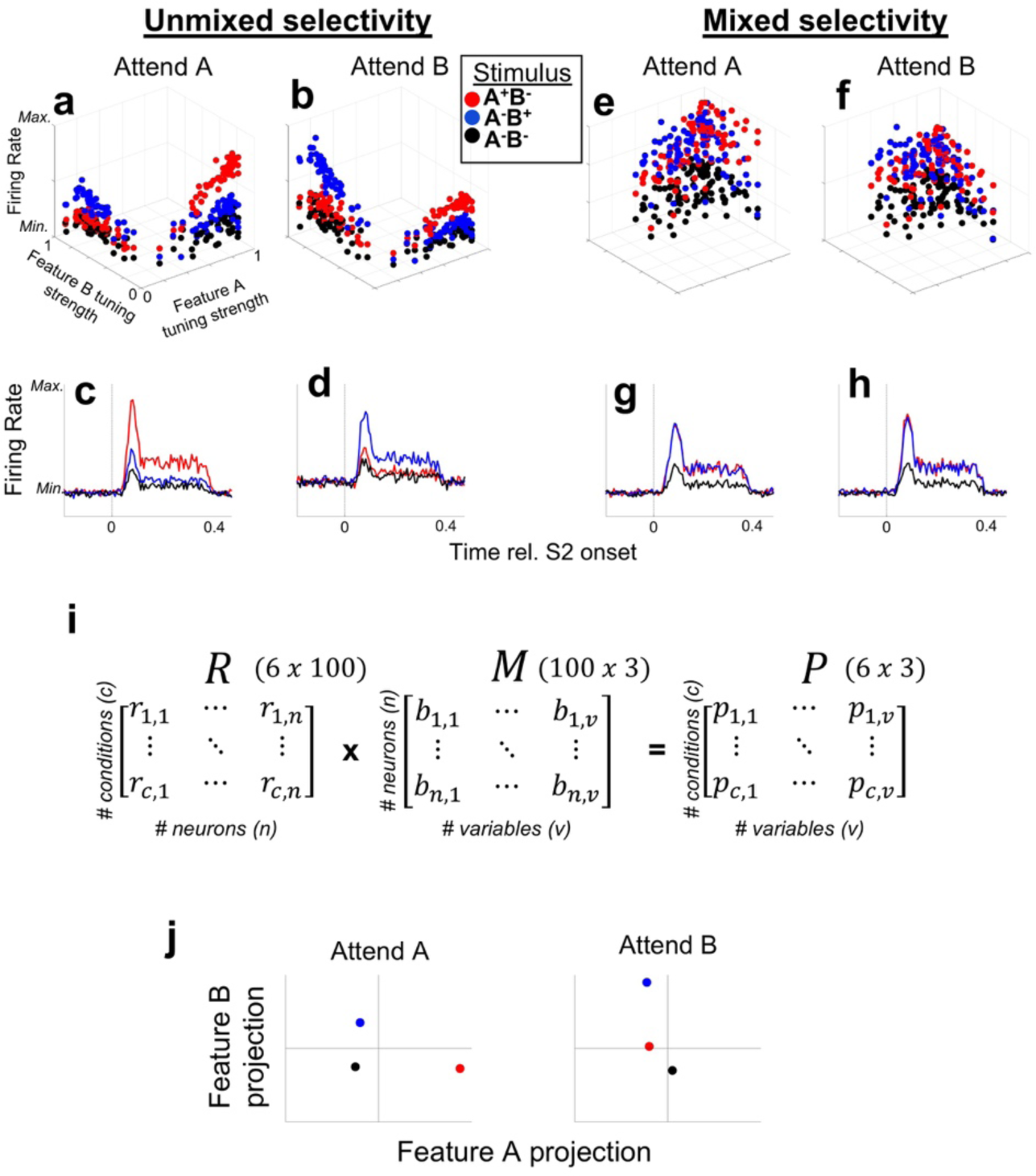
Targeted dimensionality reduction (TDR) for populations with “Mixed Selectivity”. Contrasted are two simulated populations of neurons (**a-d**, Unmixed Selectivity and **e-f**, Mixed Selectivity), each exhibiting the same Feature-Specific Gain (FSG) parameter on single neurons’ firing rates. With Unmixed Selectivity, FSG results in a simple marginalization of population firing rate that enhances the response to the attended feature (e.g., firing rate responses to Feature A [stimulus A+B-red dots and red PSTHs] exceed those of the Feature B (A-B+) and null (A-B-) stimuli when Feature A is attended (**a,c**), and vice versa when Feature B is attended (**b,d**)). **(e-h)** However, when neurons exhibit Mixed Selectivity for Features A and B (i.e. they exhibit significant tuning strength for both features), the same modeled FSG parameter does not yield an attention-related increase in population-averaged firing rate responses to the attended feature (e.g. responses to A+B- and A-B+ are roughly equal across attention conditions). However, using TDR to de-mix population response outputs **(i)** can yield population-level responses that reflect a strong enhancement of the representation of the attended vs. unattended feature. The matrix *M,* consisting of orthogonalized regression coefficients (the β in eq. 2) for each neuron, transforms the neural data in *R* to a set of 3-dimensional coordinates, (e.g., in our case the feature variables AM, ΔBW and context), for each experimental condition in matrix *P*. **(j)** A hypothetical TDR-estimated response (in arbitrary units of projection magnitude) for a population similar to that depicted in **e-h** is illustrated in **(j)** for the three stimuli (A-B-, A+B- and A-B+) for each attention condition (Attend A and Attend B).

However, if neurons’ feature tuning exhibits mixed selectivity, i.e. non-zero tuning to both features, the same FSG mechanism fails to meaningfully segregate the population-averaged firing rates (Fig. 3e-h). Thus, when single neurons exhibit mixed selectivity in their tuning to different sensory features, FSG as a mechanism of attention exhibits no average attentional enhancement (Fig. 3g,h). However, via TDR, we can find the data transformations that yield “de-mixed” representations of task variables via projections of neural population activity onto task-specific axes. In Figure 3i, we show how a full data matrix, *R* (of *z*-scored firing rates) can be transformed to a matrix *P* of uncorrelated projections via multiplication by an orthogonalized regression coefficient matrix, *M* (see next paragraph for detailed description). Thus, each of the six experimental conditions illustrated here (*c* = 3 stimulus conditions x 2 attentional; rows of *R*, and rows of *P*) yields a unique population response (projection) represented in matrix *P*. Therefore, each row of *P* represents the population response as single point in 3-dimensional space, where each dimension corresponds to a task variable (e.g. *ß*A *ß*B and *ß*context, comprising the columns in matrix *M* and *P*). This process is analogous to re-weighting each neurons’ outputs at the downstream synaptic level. Each axis in Figure 3j thus could represent the activity of a single hypothetical downstream neuron receiving its inputs from a neural population similar to that depicted in Figure 3e-h, the input weights of which are selectively shaped to maximize the unique encoding of each task variable; for clarity, we display two 2-dimensional spaces (one for each attention condition) where the dimensions correspond to the variables Feature A (x axis) and Feature B (y axis). Whereas population-averaged single-neuron firing rates fail to reveal an effect of FSG, projections onto the stimulus-variable axes may reveal a strong effect of attention: in the Attend A space, the population response to the A+B-stimulus (red) projects much farther along the Feature A axis than in the Attend B space. Likewise, the population response to the A-B+ stimulus (blue) projects farther along the Feature B axis during the Attend B space than in the Attend A space.

To implement TDR, the data are represented in matrix *R*, which contains one column for each neuron and one row for each condition. Note, in the present study we refer to “conditions” as distinct combinations of AM, ΔBW and context, such that we have 26 total experimental conditions (13 AM*ΔBW combinations across two attention conditions). We refer to “variables” as the three dimensions along which all conditions vary, namely the two sound features (AM, ΔBW) and context (attention). The matrix M consists of the set of regression coefficients calculated for each variable in Equation 2, orthogonalized via orthogonal triangular decomposition (‘qr’ command in MATLAB) to obtain the matrix. *M* is thus a set of uncorrelated variable coefficient vectors that can be used to project our data onto orthogonal axes corresponding to task variables in the form of matrix *P*. We illustrate the process of TDR on our data in Figure 4. We have uploaded Matlab code demonstrating the analysis steps for running TDR to github.com/joshddowner.

**Figure 4.**
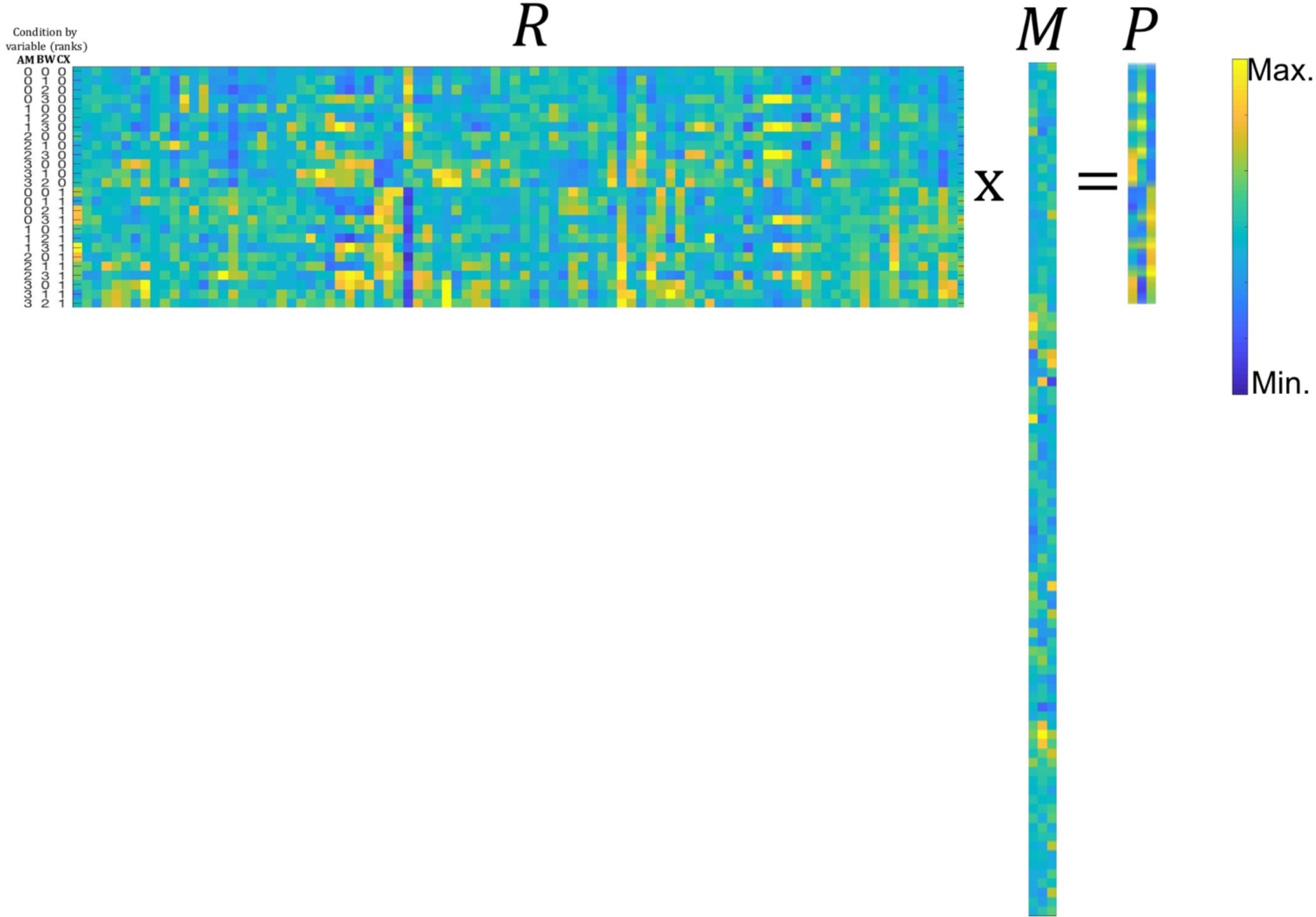
Concrete description of matrices *R, M* and *P* as expressed in our data. Across matrices, the color scheme parula represents the values (firing rate in *R*, coefficient value in *M*, and projection magnitude in *P*) of the entries in each matrix (maxima: yellow; minima: blue). The matrix *R* is 26*92, where each row corresponds to a condition (all 26 combinations of 4 AM depths, 4 bandwidths, and two attend conditions) and each column to a neuron (92 neurons). The matrix *M* is 92*3 where each row corresponds to a neuron and each column to a vector of orthogonalized variable coefficients (βAM, βBW and βcontext from equation 2). Here, the columns of *M* are (1) AM coefficients, (2) BW coefficients and (3) context coefficients. Multiplying *R x M* yields the matrix *P* in which each row corresponds to one of the 26 conditions and each column to a variable, or axis. Each value of *P* represents the projection of the population response to a condition along a given axis. For instance, column 1 of *P* exhibits increasing values as the rank value of AM increases (see condition labels to the left of matrix *R*). The color bar on the right applies to all matrices and indicates values from minimum (blue) to maximum (yellow) for firing rate (matrix R), coefficient (matrix M) and projection (matrix P).

### Simulating neural population activity

Since many of our neurons were recorded during different sessions, we used a bootstrapping approach to estimate the mean and variance of population projections. Namely, we constructed neural populations of size *n* = 92 neurons by randomly sampling, with replacement, from our set of 92 sampled A1 neurons. We similarly simulated trials by re-sampling (with replacement) from the distribution of spike counts of each neuron in each of the 26 experimental conditions (10 trials per condition).

An important consideration with this approach is that, by combining neurons across experimental sessions, and by randomly re-sampling trials, we fail to approximate correlated and intrinsic variability between and within neurons, respectively. Therefore, the matrix of simulated spike counts within a condition is transformed to introduce noise correlations and neuron-intrinsic variability that match that observed in the data. We determined the desired noise correlation value between a given pair by measuring the noise correlations between the simultaneously recorded pairs of neurons in the data (n = 434 pairs) and fitting a linear regression to those noise correlation values to determine the impact of AM and ΔBW tuning correlation, joint firing rate and attention condition on noise correlations. We determined the desired neuron-intrinsic variability by calculating the Fano factor (variance/mean rate) for each neuron and scaling its simulated spike count distribution to match the observed Fano factor. The methods for introducing realistic noise correlations and Fano factor in simulated neural populations is detailed in Shadlen and Newsome (Shadlen and Newsome, 1998) and Downer et al. (Downer et al., 2017).

We determined the sensitivity for each feature in each simulated population by calculating the ROC area (as described for single neurons) by comparing distributions of simulated trial-by-trial projections onto each sound feature axis (as opposed to distributions of trial-by-trial spike count distributions). The simulated projections in a given trial were calculated as in Figure 3i, where *R* is a matrix of condition-normalized (*z*-scored across conditions) spike counts (columns) across conditions (rows), *M* is the matrix of orthogonalized regression coefficients and *P* constitutes the 3-dimensional projection for each of the 26 experimental conditions. We calculated decoding accuracy (ROC area) for AM and BW by comparing the distribution of projections of AM>0 vs. AM0 stimuli and BW>0 vs. BW0 stimuli, respectively, along each feature’s axis. A linear support vector machine was fit to a training set and performance evaluated on a test set of withheld data. We calculated choice-related activity by comparing projection distributions along each axis for “yes” and “no” response trials, and by using the same support vector machine algorithm to discover the linear plane of separation between the 3-dimensional (AM*BW*Context) projections for “Yes” and “No” trials. Importantly, we only analyzed choice-related population activity for stimuli with roughly equal numbers of “yes” and “no” responses –namely, those stimuli near perceptual threshold, where subjects made at least 5 correct and incorrect judgements, similar to Niwa et al. (Niwa et al., 2012a). This provision protects against biases arising from re-sampling from distributions with few values, for which mean and variance estimates are unreliable.

In order to evaluate the extent to which findings of population-level activity deviate from expected effects of pooling across groups of neurons, we constructed 1000 “surrogate” populations with intact single-neuron tuning but decimated higher-order correlations among them (Elsayed and Cunningham, 2017). This method constructs surrogate populations by shuffling the original data labels, but then recovering (to the extent possible) the marginal and covariance features of the original data by transforming the shuffled data using a “readout” matrix. Thus, the final surrogate populations contain modeled neurons that closely approximate the neurons in the original data, but with higher-order correlations abolished. We calculated sensory sensitivity (ROC area) and choice-related activity precisely as we did with the intact data set, across 1000 surrogate populations. These distributions produced by the surrogate populations provide benchmarks for the expected results given effects accountable by single-neuron features without complex interactions amongst them.

## RESULTS

### Monkeys successfully perform the feature selective attention task

Within each feature-attention (context) block, we quantified the degree to which a change in each feature influenced subjects’ probability of a “Yes” response using a binomial logistic regression (see Methods section). Each subject performed the task above chance, as evidenced by a greater influence of the attended feature vs. the unattended feature on the probability of a “Yes” response (Figure 2; 47 total sessions, *p = 8.98e^-8^*; sign rank test, collapsed across monkeys).

### A1 neurons exhibit mixed selectivity for sensory features and task context

We quantified encoding of behavioral (context) and acoustic (AM and BW) variables on the firing rates of recorded A1 neurons (n = 92) using multiple linear regression (Equation 2). For each neuron, we calculated a set of coefficients describing the main effect of each of these three variables and all pairwise interactions (9 total coefficients per neuron).

We saw diverse encoding of task variables, as shown in several example neurons (Figure 5; each column of panels is a different neuron). The top row of Fig. 5a-c shows firing rates, collapsed across context and BW condition, separated by AM condition (red: AM>0; black: AM0= No AM). The middle row shows firing rates separated by BW condition, and the bottom row shows firing rates separated by context. The neuron in 3a encodes BW, shown by a decreased firing rate for BW>0 conditions relative to BW0 conditions, while the neurons in 5b and 5c encode both sound features as well as context, albeit with different directions of firing rate changes (e.g. they both encode AM, but 5b exhibits an increased firing rate for AM while 5c exhibits a decreased firing rate). Figure 5d-f shows the time-course of the coefficient values calculated for each variable by the neurons in 5a-c, respectively. Taken together, these 3 example neurons illustrate that single A1 neurons exhibit diverse encoding of variables relevant for this task, often encoding context as well as both sound features. Following from this finding is the possibility that because A1 firing rates simultaneously encode multiple independent variables, single-neuron firing rates may provide a poor code for any given variable on a trial by trial basis, i.e., that the unweighted firing rates of individual A1 neurons do not unambiguously encode the variables necessary for task performance. Therefore, marginalizing across single neurons by unweighted averaging yields an ambiguous representation. A summary across all single neurons is shown in Figure 6.

**Figure 5.**
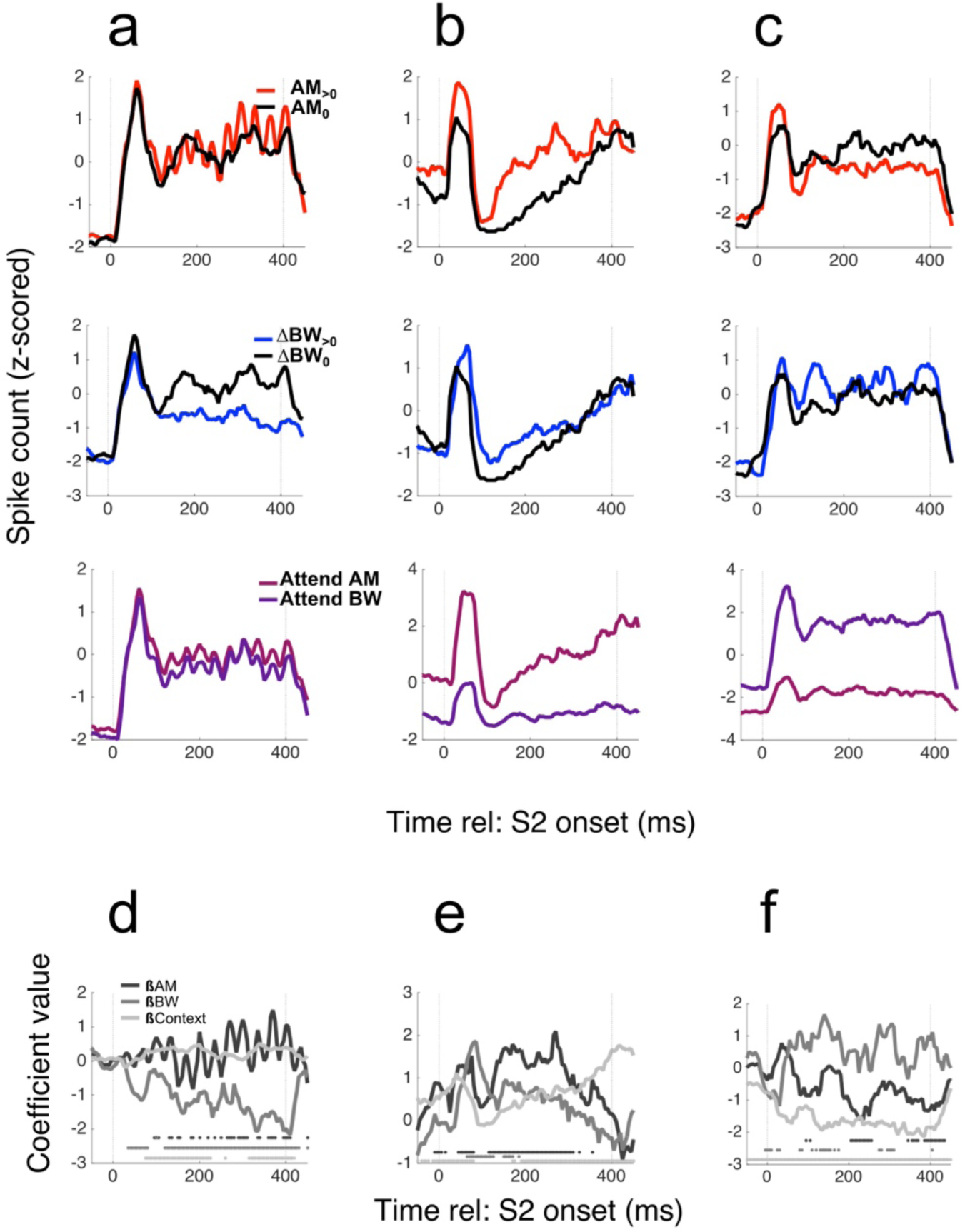
A1 neurons exhibit mixed selectivity for sensory and task variables. Example PSTHs for 3 neurons are shown in columns (**a**), (**b**) and (**c**), with average firing rates collapsed across irrelevant dimensions to reveal AM (top row), BW (middle) and Context (bottom) encoding. The neuron in (a) exhibits no AM- or Context-dependent changes in sustained firing rate, but decreases its firing rate when ΔBW>0. The neuron in (**b**) increases its rate for both AM>0 and ΔBW>0 (red PSTH and blue PSTH above black PSTH, respectively) and has an overall higher firing rate during the Attend AM context (pink PSTH above purple PSTH). The neuron in (**c**) decreases rate for AM>0, increases rate for ΔBW>0 and prefers the Attend BW condition. We calculated coefficients for each variable over a sliding window (0 means that variable does not contribute to firing rate), and these time-varying coefficient trajectories are shown in (**df**) for the neurons in (a-c), respectively. The greyscale markers along the ‘x’ axis in the bottom of each panel indicate significance of each coefficient within the 50-ms time bin in which the coefficient was calculated.

**Figure 6.**
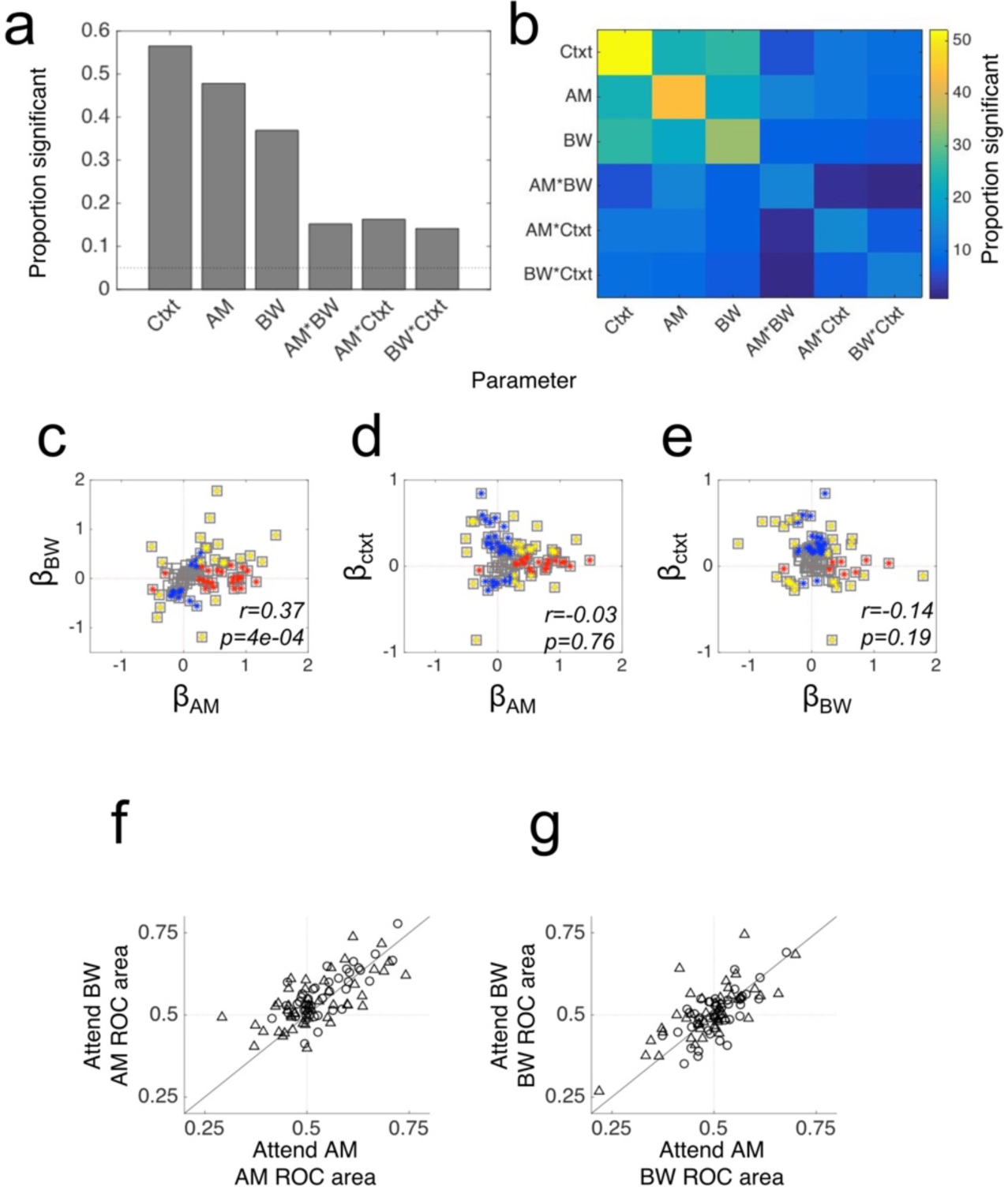
A1 neurons robustly encode Context, AM and BW. (**a**) The histogram showing the proportion of neurons with significant Context, AM and BW coefficients reveals that each variable modulates the firing rate of many A1 neurons. Importantly, interactions between the variables are relatively rare, suggesting that each variable tends to make an independent, linear contribution to neurons’ firing rates. Most obviously, the paucity of cells with significant Context*AM or Context*BW interaction terms suggests that sound feature tuning in A1 does not vary with attention condition. (**b**) Many neurons that encode one variable also encode at least one other. For instance, more than half of those cells that encode context also encode either AM or BW. (**c**) There is a positive relationship between AM and BW encoding, such that the single neuron firing rates will tend to fail to disambiguate between AM and BW. (**d** & **e**). Context exhibits no significant relationship between AM or BW tuning, revealing that the feature-specific gain model is inconsistent with our data. In (**c-e**), red markers indicate a significant coefficient for the variable on the ‘x’ axis, blue markers indicate a significant coefficient for the variable on the ‘y’ axis and gold markers indicate significance for both variables. **(f)** Single-neuron feature sensitivity is consistent across attention conditions. Single-neuron AM sensitivity, as measured with ROC area (AM ROC area) does not change between conditions, on average. **(g)** Same as (f), but for BW ROC area.

Context had a significant main effect in more neurons than either acoustic feature, as measured by the proportion of neurons with significant context coefficients (Fig. 6a). Significant interaction effects were less common, suggesting that attention does not tend to change A1 neurons’ feature tuning. This can be seen clearly in Figure 6b as well: the color map of proportions of significant coefficients exhibits a markedly higher proportion of main effects (main diagonal) than interactions. This contrasts with findings from tasks comparing passive listening and active auditory behaviors on A1 tuning: across many studies, active task engagement result in large changes in A1 tuning (David, 2018). These studies have been interpreted as evidence that A1 single-neuron tuning tracks behavioral demands, namely that tuning is enhanced when animals are actively engaged with sounds (Niwa et al., 2012b).

Figure 6c provides a possible reason for the absence of attention-related changes in A1 neuron tuning. We plot the value of AM coefficients against the value of BW coefficients and find that a majority of neurons have similar AM and BW tuning (as measured by the sign of the coefficient (Spearman correlation, r = 0.37, p < 0.001). In other words, A1 neurons rarely uniquely encode one sound feature or the other because feature tuning co-varies at the single-neuron level (similar to Fig. 3e-h). This prevents single-neuron tuning from providing a robust basis for attentional perceptual enhancement. We also fail to find a significant relationship between context encoding and feature tuning, as shown in Figure 6d & e. A common finding in the attention literature is that feature attention leads to “feature-specific gain”, whereby neurons tuned to the attended feature exhibit an increased firing rate (Martinez-Trujillo and Treue, 2004). An FSG effect in these data would manifest as a positive relationship between *β* AM and *β* Ctxt and a negative relationship between and *β* BW and *β* Ctxt. On the contrary, our analyses reveal that attention has no net effect on A1 neurons’ rate-based feature tuning. Thus, neither attention-related changes in tuning functions, nor feature-specific gain effects are noticeably present in the firing rates across the individual neurons we observed.

### Average A1 neurons’ feature sensitivity is constant across attention conditions

In the analyses presented in Figure 6a-e, we calculated coefficients across the entire experiment and used sound feature × context interaction terms as a measure of attentional modulation of A1 neuron tuning. We next measured the average effects of attention on single-neuron feature decoding accuracy by calculating ROC area for each feature in each attention condition for each neuron (see Methods). Comparing ROC area for both features across conditions reveals no attentional modulation of single-neuron feature sensitivity (Figure 6f,g; results shown are collapsed across all AM and BW levels). In 6f, the AM ROC area is shown for the Attend AM condition on the x axis and the Attend BW condition on the y axis. Though we do observe some scatter around the unity line, neurons increase or decrease their AM ROC area with equal likelihood, (sign rank test, p = 0.74). In 6g we show the BW ROC areas across attention conditions and, again, find no average effect of attention on this single-neuron metric of feature tuning (p = 0.73). Therefore, when comparing feature tuning across the more rigorous demands of different feature attention conditions, we do not observe tuning changes that have been found in several prior studies comparing A1 tuning between passive vs. active conditions and between attend-toward vs. attend-away conditions (Niwa et al., 2012b; Schwartz and David, 2017; von Trapp et al., 2016).

### Targeted dimensionality reduction analysis reveals attentional enhancement of sensory encoding

Given the complexity of the task, we reasoned that a plausible strategy for adaptive feature selection might be a distributed population code. Population codes can allow unambiguous and simultaneous encoding of many variables by distributing the task-relevant signals across single neurons (Fusi et al., 2016). Thus, during population encoding of complex auditory scenes involving multiple relevant variables, single-neuron firing rates may provide a complicated snapshot of the overall operations of A1. We therefore modeled A1 population coding by subsampling from our set of single neurons, and projected population vector activity (Georgopoulos et al., 1986) onto axes corresponding to AM and BW encoding using Targeted Dimensionality Reduction (TDR) (cf Mante et al 2013; see Methods and Figures 2 and S2). Such approaches have been employed in studies of higher-order cortex to disambiguate the often-heterogeneous single neuron signals observed there (Machens et al., 2010; Mante et al., 2013; Rigotti et al., 2013).

We provide a description of population responses across experimental conditions in Figure 7. Here, population-averaged firing rates (averaged over the entire 400-ms S2 presentation epoch, Fig. 1a) are displayed in 4x4 grids, where each element represents firing rate averaged over a sub-group of neurons sorted by *β* AM (x-axis) and *β* BW _(y-axis)._ A hypothetical population response in Fig 7a shows a case in which neurons with positive *β* AM and positive *β* BW fire above their mean rates, whereas neurons with negative *β* AM and *β* BW fire below their mean rates. Our data displayed in this manner qualitatively reveal effects of stimulus feature and task context (Fig. 7b,c). Each panel in Fig. 7b & c shows the population response (as in Fig. 7a) for a given stimulus combination of AM and ΔBW (for example, the lower left panel in Fig. 7b is the population response in attend-AM to the ΔBW0-AM3 stimulus).

**Figure 7.**
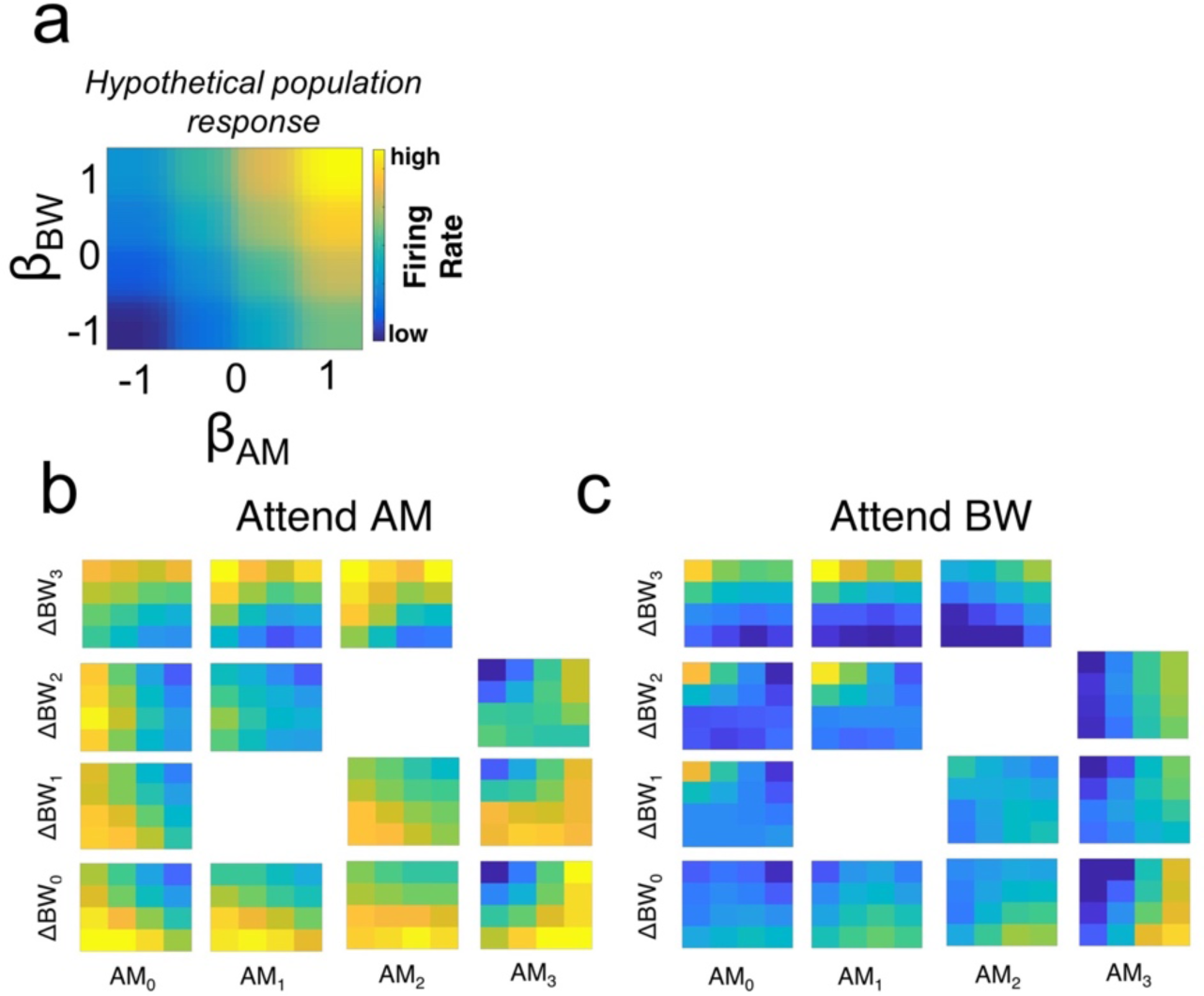
A population-level analysis for feature encoding in A1. (**a**) We provide a schematic illustration of our population-level analysis approach. In the grid in (a) is a hypothetical population response pattern, with population firing rate organized by ßAM and ßBW. For this hypothetical response, the group of neurons with positive ßAM and ßBW exhibit high firing rates. (**b**) In our data, population response patterns differ across the stimulus set (population response grids arranged as the stimulus matrix in 1b), with clear differences in population responses related to increasing AM and BW levels. Moreover, comparing (**c**) to (**b**) reveals substantial differences in population responses related to attention.

By and large, increases in AM level result in population activity patterns that are increasingly biased toward high spike counts among neurons with positive *β*_AM_ and increases in ΔBW level result in patterns with high spike counts among neurons with positive *β*_BW_. Moreover, comparing population activity patterns for a given stimulus across attention conditions hints at complex changes in firing rate based on both *β*_AM_ and *β*_BW_. For instance, for AM0-ΔBW2, the population activity pattern during Attend AM seems to exhibit high firing rates for neurons with negative *β*_AM_, regardless of *β*_BW_. However, during Attend BW, the population activity exhibits low firing rates for all neurons except for those with negative *β*_AM_ and positive *β*_BW_. Taken together, these panels provide a visual intuition for how stimuli are encoded in a distributed manner across the A1 population and how attention qualitatively affects this encoding.

Projecting population activity into a low-dimensional subspace via TDR provides a compact way of quantifying population response patterns (Figure 8). We focus here on stimulus-related projections. In Figure 8a, 8 distinct, idealized population responses show where each population response falls on an AM axis projection versus BW axis projection plot. So the population response in the upper right of Fig. 8A, would have a value near 1 for both its AM and BW axes projections, and thus would be a dot in the upper right for panels 8b-d. In order to project our data onto these coordinates, we constructed pseudo-populations by sampling from our pool of recorded neurons and simulating trial-by-trial population responses, which are projected onto these 2D coordinates via a linear combination between a data matrix of spike counts (*R*) and a coefficients matrix (*M*) (see Fig. 3i,j and Methods). Projections in the Attend AM and Attend BW conditions for each stimulus are shown in Figure 8b and c, respectively; projections averaged over simulated trials are shown for each stimulus condition (see inset for color code). Stimulus encoding is evidenced by increasing AM level corresponding to greater projections along the AM axis, and increasing ΔBW levels correspond to greater projections along the BW axis (e.g. AM3 and BW3 stimuli project farthest along the AM and BW axes, respectively). In addition, the topography of stimulus responses clearly differs between attention conditions. Thus, whereas single-neuron firing rates fail to disambiguate between AM and BW, populations comprising these same neurons successfully encode each feature when their outputs are re-weighted using TDR.

**Figure 8.**
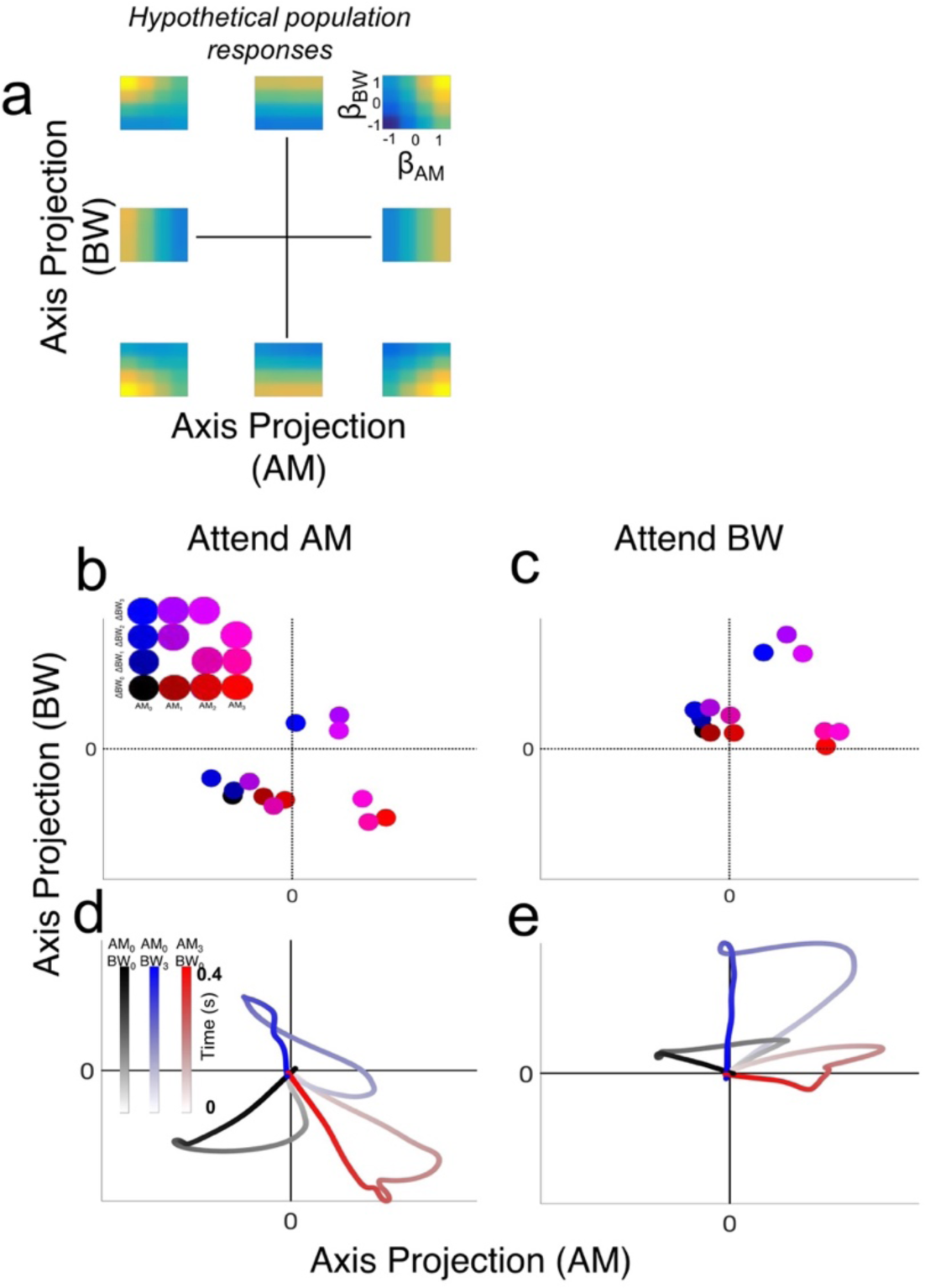
Neural activity projected into sensory-defined subspace reveals sound feature encoding. (**a**) Our approach involves projecting responses of populations of neurons into lower-dimensional *condition* subspace defined by the stimulus variables (dimensionality reduced from *n neurons* to *p conditions*). The eight hypothetical responses in (a) show idealized population responses that project in corresponding eight points in the AM axis projection versus BW axis projection coordinates. (Context dimension not shown). (**b**) Color-coded stimulus responses (see inset) in the Attend AM condition are shown projected into axis-projection subspace. Large symbols represent mean responses over simulated trials. Markers representing projections along the AM axis exhibit increasing (*red*) values, whereas markers representing projections along the BW axis exhibit increasing (*blue*) values, indicating population-level encoding of sound features. In the upper right quadrant are purple markers, representing stimuli containing both features, and markers with low color values (including black) are found in the lower left quadrant. (**c**) Same as (b) but for the Attend BW condition. Axes in (b) and (c) are scaled equally. Comparing encoding subspaces across conditions reveals substantial effects of attention on sensory subspace projections. Namely, near-threshold stimuli exhibit stronger relative projections along a given feature axis when that feature is attended. (**d**) Trajectories over time through the sensory subspace for 3 representative stimuli (AM0-BW0 [black], AM0-BW3 [blue], AM3-BW0 [red]) reveal encoding time courses over the entire course of S2 presentation. Trajectories begin at the ‘x’ and ‘y’ origin, indicating no sensory evidence for either AM or BW. Directly after stimulus onset, trajectories for each stimulus follow similar paths, then diverge substantially. Trajectories reach a peak of separation, such that AM0-BW3 & AM3-BW0 projections are in opposite quadrants (upper left and lower right, respectively), both roughly orthogonal to the AM0-BW0 projection. Then, during the end of the stimulus response, trajectories return to 0. (**e**) Same as in (d) but for the Attend BW condition. Trajectories in both conditions exhibit an early, non-selective course wherein each stimulus projects to roughly the same area within the subspace. During the middle of the stimulus, trajectories reach their maximum separation, then quickly return to 0. These trajectories suggest that feature selection during sound perception evolves over time, beginning with a general detection phase, followed by a discrimination phase, then returning to a non-encoding area of the subspace quickly thereafter.

Moreover, we observe that trajectories through this subspace exhibit interesting temporal dynamics (Figure 8d,e). For clarity, only 3 stimulus response trajectories are shown (AM0BW0 [black], AM3BW0 [red], and AM0BW3 [blue]). Time is shown with lighter shading of the lines early in the stimulus gradually getting darker over time. The initial excursion for each stimulus, corresponding to the onset response immediately after time 0, appears initially ambiguous and gradually differentiates over the course of a S2 presentation. Projection trajectories appear maximally separate roughly in the middle of the stimulus, and by the end of the stimulus presentation, trajectories return to the origin of the subspace. These temporal dynamics suggest that early population responses function to detect a rapid change, irrespective of feature, whereas sustained responses elicit finer-grained feature selectivity. Differences between onset and sustained responding have been widely observed in single auditory cortical neurons (Osman et al., 2018). These findings support that the temporal dynamics of sensory neuron responses correspond to distinct encoding stages in population activity patterns.

Qualitative effects of attention are apparent when comparing the population activity patterns across Attend AM and Attend BW conditions (e.g. Fig. 7b vs. 7c, Fig. 8b,d vs. 8c,e). We next quantified to what extent population encoding of relevant vs. irrelevant stimulus features was affected by these changes. At the single-neuron firing rate level, we observe no effect of attention enhancing the representation of attended vs. unattended features (Figure 6f,g; Figure 9a). However, using signal detection analyses on TDR population responses (see Methods) we find that population projections onto 2D weighted-feature space afford better decoding accuracy for each feature when it is attended (Figure 9b). Across 100 simulated neural populations, the average ROC enhancement is 0.151 for AM (*p=2.41e-30*) and 0.091 for BW (*p=3.08e-17*).

**Figure 9.**
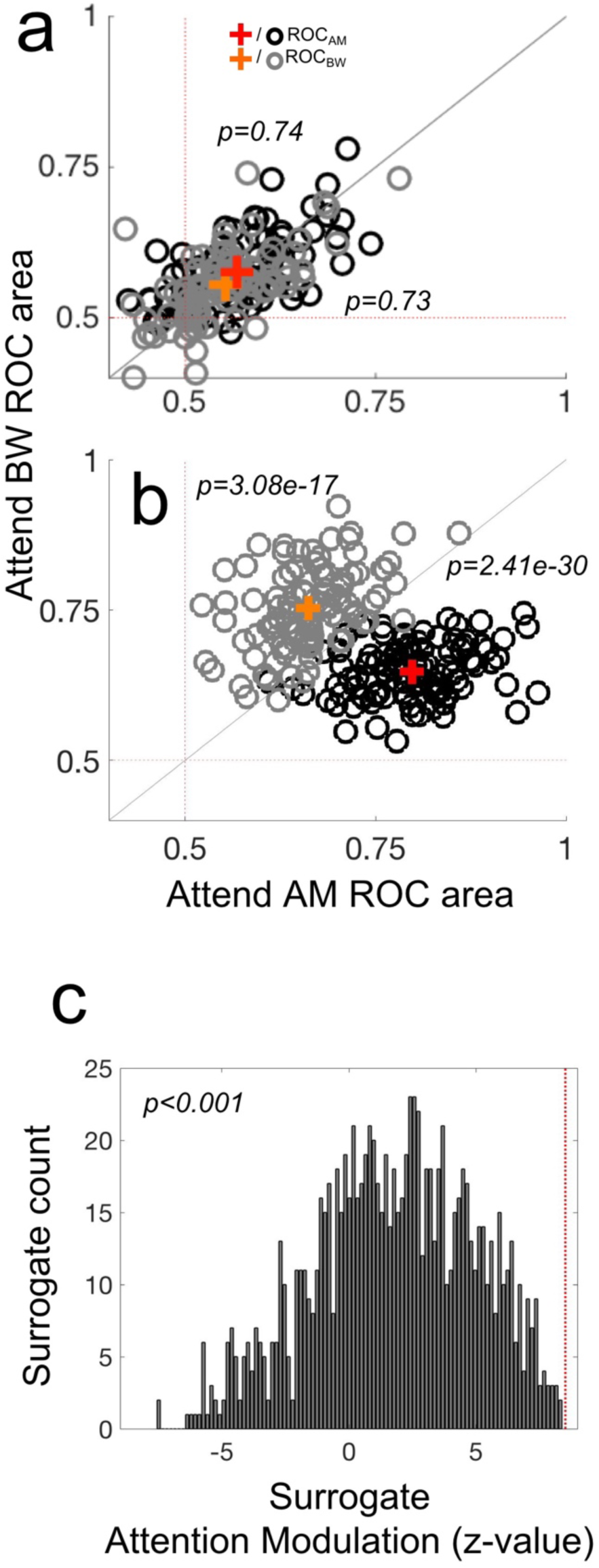
Population, not single neuron, representations sharpen with attention. (**a**) Single-neuron ROC areas (similar to Figure 4f,g) are compared between the Attend AM (x-axis) and Attend BW (y-axis) conditions. Black ‘**o**’ markers represent single neuron average AM ROC values, with the red ‘**+**’ representing the mean AM ROC value across the population. Grey ‘**o**’ and orange ‘**+**’ represent single-neuron and mean BW ROC values, respectively. AM ROC and BW ROC distributions each lie along the unity line, indicating no average effect of attention (*p = 0.73* and *0.74*, respectively). (**b**) Simulated population ROC areas (one black and one grey marker for each population of size *n* = 100 neurons) differ significantly between attention conditions, such that the average AM ROC area is 0.151 higher during Attend AM than Attend BW (red ‘**+**’; *p = 2.41e -30*) and the average BW ROC areas is 0.06 higher during Attend BW than Attend AM (orange ‘**+**’; *p = 3.08e-17*). (**c**) The distribution of Attention Modulation values (the z-scored increase in ROC area for each feature when it is attended vs. ignored) for 1000 surrogate populations is compared against the observed Attention Modulation in the neural data (vertical red line). Though surrogate populations, on average, exhibit a positive Attention Modulation (as evidence by the peak of the distribution well above 0), the Attention Modulation observed in the neural data exceeds that of any given surrogate population. This indicates high confidence that the finding in (b) constitutes a synergistic effect of attention on neural coding, above and beyond what would be expected if a weak attentional enhancement was present in a majority of individual neurons.

This dissociation between single-neuron and population attentional enhancement suggests a prioritized role for population-level representations in primary sensory cortex during feature selective attention. Importantly, the difference in sensory sensitivity effects between single neurons and populations cannot be attributed to greater analytical power obtained by selective pooling over many neurons (Figure 9c).

Comparing the attentional modulation index (z-score of ROC difference of attended relative to unattended feature) of the neural data (9c, vertical red line) to that of 1000 surrogate populations, we find no overlap between our observed attentional enhancement and that expected by the simpler single-neuron and pairwise covariance properties. This suggests that our observed population enhancement of sound feature encoding constitutes an emergent effect, relying on higher-order correlations among A1 neurons.

Finally, we analyzed how correct vs. incorrect performance accounts for variance in trial-by-trial projections within the stimuli near perceptual threshold (AM1-ΔBW0 and AM0-ΔBW1, Fig. 10). In order to quantify these effects, we calculated the ROC area between correct and incorrect response distributions, a metric commonly referred to as “Choice Probability” (CP) (Britten et al., 1992). In Fig. 10a-c, we show the CP associated with variance along each task variable axis. For each single-axis projection, we find a significant difference from 0.5 in CP across 100 simulated neural populations. Moreover, we also assessed CP for projections into the 3-dimensional (AM*BW*Context) subspace using a support vector machine algorithm to define a plane that best separates correct from incorrect trials’ projections (Fig. 10d). This 3D-projection exhibited especially large CP values relative to projections onto the individual axes. Thus, choice-related activity accounts for some unique proportion of variance in the population data. Taken together with the attentional enhancements, the observation of performance-related population-level activity in A1 supports the notion that neurons in A1 act cooperatively to support auditory perception and decision-making.

**Figure 10.**
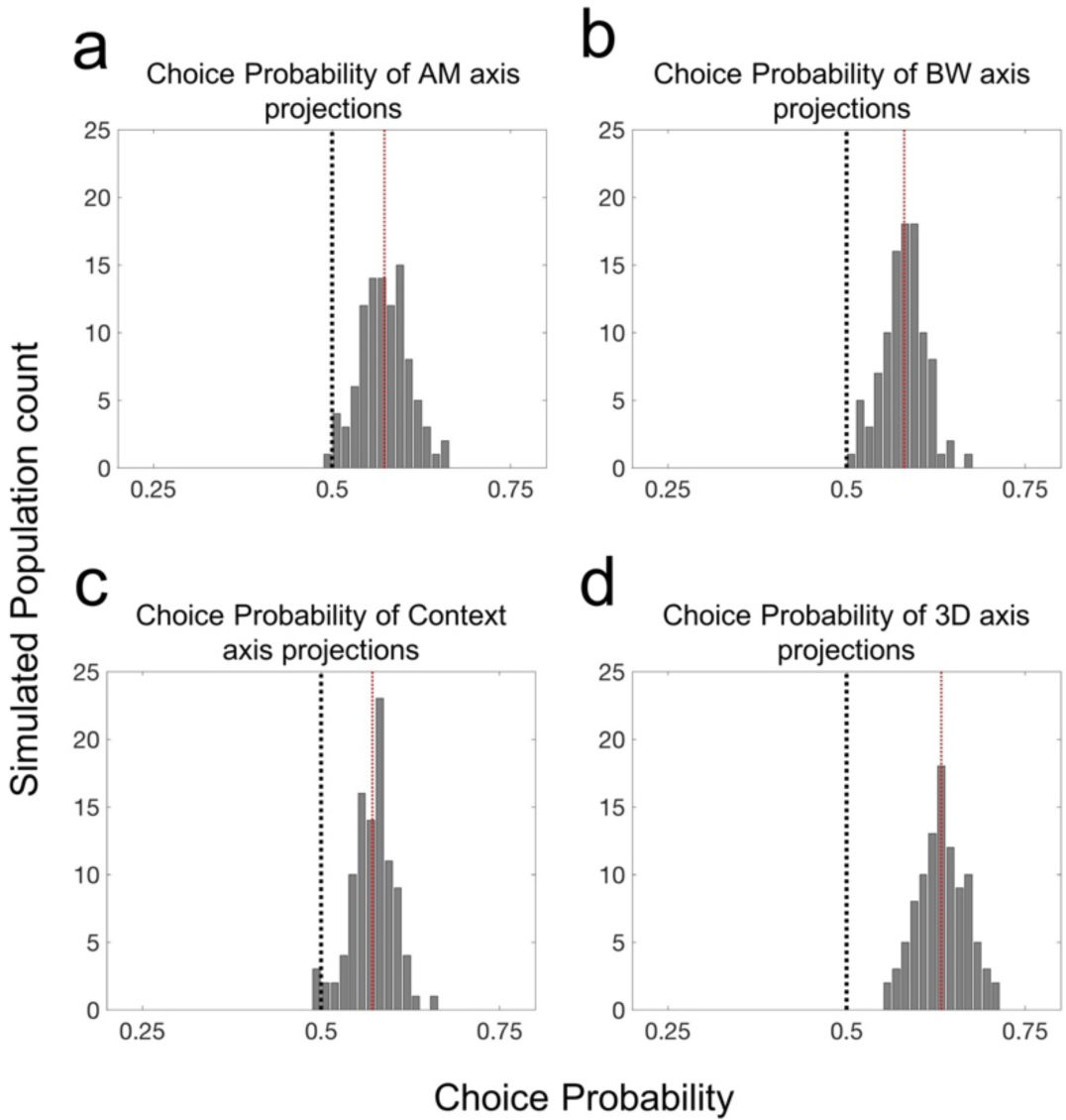
Population subspace projections correlate with trial-by-trial performance. (**a**) The distribution of choice probability (CP) values across 100 simulated populations reveals a significant relationship between AM-axis projections and task performance, as evidenced by a mean CP across simulations of 0.573 (*p=1.90e-39*). Likewise, (**b**) & (**c**) reveal that projections along the BW and Context axes also correlate with task performance, with mean CP values of 0.581 and 0.572, respectively (*p=4.82e-47* and *1.37e-42*). (**d**) Projections of population activity into the full 3-dimensional task variable subspace (AM*BW*Context) were very reliably performance-related, as evidenced by the entire distribution of CP values across simulated populations lying above 0.5, with a mean of 0.633 (*p=1.12e-61*). Each significance test was a single-distribution t-test with a null hypothesis of CP=0.5.

## DISCUSSION

We provide evidence that population-level encoding of sound features is more informative than single-neuron encoding. In general, averaging across many observations yields more reliable estimates of observed properties, and therefore the finding that groups of neurons outperform single-neuron sensory encoding perhaps seems unsurprising. However, many phenomena in neural circuits violate basic parametric statistical assumptions (e.g. saturation, thresholding, etc.), therefore disallowing simple averaging or pooling across neurons (Goldman et al., 2001), while novel analyses afford the opportunity to disambiguate the expected population-level enhancements from those observed (Elsayed and Cunningham, 2017). We find that single-neuron representations are insufficient to explain attention- and choice-related activity in neural populations in A1. Instead, a population-level description is required to link sensory cortical activity and perception in this primary sensory cortical area.

In the last several years, multiple studies of A1 have suggested the importance of population-level representations. Studies have found that population-level analyses provide clear links between A1 activity and behavior, whereas single neurons may not (Bagur et al., 2018; Christison-Lagay et al., 2017; See et al., 2018; Yao and Sanes, 2018). A critical question has remained unanswered, however: To what extent are these population-level findings an expected by-product of marginalizing across neurons (Sasaki et al., 2017)? For instance, in comparisons between active and passive states, in which only a single sensory feature matters for task performance, single neurons will likely resemble noisy versions of population representations, simply by virtue of there being very few stimulus parameters influencing single neurons. Studies of high-dimensional (e.g. multi-feature) sensory representation in single A1 neurons during passive sound presentation have shown that single A1 neurons do indeed exhibit high dimensionality (O’Connor et al., 2005, 2010; Sadagopan and Wang, 2009; Sloas et al., 2016) which can provide unambiguous population coding beyond that provided by any single neuron (Bizley et al., 2010). Importantly, in the present report, sufficiently high dimensionality of the sensory and behavioral variables of the task allows for a direct test of high-dimensional representation in single neurons and whether single-neuron representation suffices to explain population findings related to perception. We find that single neurons represent task variables in substantially “mixed” fashion, such that single-neuron firing rates provide highly ambiguous information (Fusi et al., 2016). Only at the level of population encoding does A1 activity correlate with perception, both in terms of attentional enhancement of the encoding of the attended feature and in terms of explaining variance related to subjects’ perceptual reports. Such mixed selectivity/population-level primacy tends to be associated with association and pre-frontal cortices. Our findings therefore highlight the importance of studying primary sensory cortical neurons in sufficiently complex experimental conditions to uncover their basic properties.

An important consideration of the present report is our exclusive use of firing rate, as opposed to spike-timing measures of neural activity. A1 neurons across species encode temporal modulations with phase-locked spikes (Johnson et al., 2012; Lu et al., 2001; Malone et al., 2007; Yin et al., 2011) and previous studies have reported significant (albeit weak) effects of task engagement on spike timing (Niwa et al., 2015; Yin et al., 2014), though in A1 these effects are modest relative to those observed for firing rate codes. In our data, we find a substantial proportion of neurons that phase-lock to the temporal envelope of sounds. However, we find no evidence of attention-related effects on phase locking (Figure 11), suggesting that phase-locking may not be relevant for feature-selective attention in the context of this task in A1. It is important to note that the design of our task is such that, even when attending to the spectral feature, many target sounds are temporally modulated as well, and therefore phase-locking can signal the presence of a target in both attention conditions. Thus, the present task may not sufficiently challenge the auditory system to drive attention-related changes in the temporal dynamics of A1 neurons’ spiking. It remains an open question whether attention affects temporal coding in other tasks or in other auditory areas, since firing rate dynamics provide a powerful, explicit code for temporal sound features that can help to disambiguate neural representations when firing rates fail to do so. For instance, a task in which subjects must discriminate rather than detect temporal modulations may very well require attentional modulation of spike timing. To date, few studies have measured A1 activity during a temporal modulation discrimination task. Future studies may indeed find significant attention-related changes in temporal response dynamics in A1 or other auditory structures.

**Figure 11.**
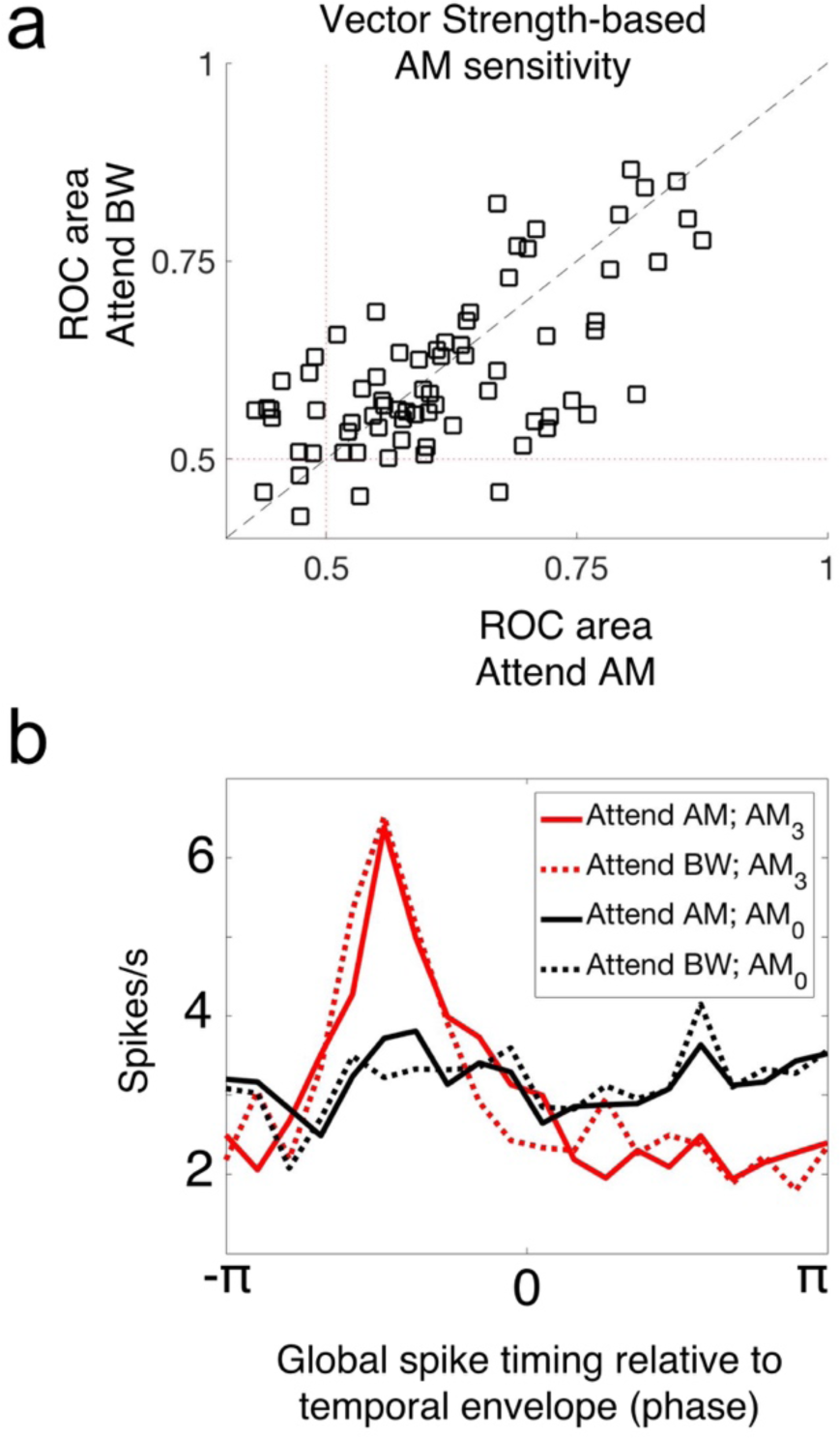
No evidence of attention-related changes in spike-timing-based sound encoding. **(a)** We plot the vector strength-based neural sensitivity (ROC area) of each A1 neuron across attention conditions (Attend AM, x axis; Attend BW, y axis). There is no average difference in ROC area between conditions (sign rank test, p=0.8849). **(b)** We analyzed the tendency for neurons to fire spikes in the same phase relative to the temporal modulation. Global spike timing in response to the fully temporally modulated stimulus (*AM3, red*) shows a prominent peak relative to the unmodulated stimulus (*AM0, black*), indicating that A1 neurons tend to fire synchronous spikes relative to temporal modulation. We observe no effect of attention on this property of A1 neurons (solid lines vs. dashed lines). These results indicate a lack of attentional modulation on the response dynamics of A1 neurons.

Some types of early sensory neural representations are inherently high-dimensional – for instance, the perception of odorants. Traditionally, primary sensory areas A1, S1 and V1 have been described in terms of a low-dimensional property, namely the placement of the stimulus on the sensory epithelium. However, no such “place” code exists for olfaction, since combinations of odorants comprise the ‘adequate stimulus’ for olfaction and a 2-dimensional sensory receptor surface can’t achieve a simple low-dimensional mapping of this combination space. Like olfaction, auditory, somatic and visual perception also involve the combinations of many basic features.

Therefore, when probed in high-dimensional conditions, these primary sensory cortical fields may reflect a similar mechanism of representation as that observed in chemosensory areas. Therefore, in contrast to the classic descriptions, primary sensory cortices may be best described in terms of their complex, combinatorial, population-level representations.

## Acknowledgements

This work was funded by NIH NIDCD grant 02514 (MLS), NSF GRFP fellowship 1148897 (JD) and an ARCS Foundation Fellowship (JD). We thank the members of the Shenoy lab at Stanford for helpful feedback.

